# The Plant PTM Viewer, a central resource exploring plant protein modifications. From site-seeing to protein function

**DOI:** 10.1101/415802

**Authors:** Patrick Willems, Alison Horne, Sofie Goormachtig, Ive De Smet, Alexander Botzki, Frank Van Breusegem, Kris Gevaert

## Abstract

Posttranslational modifications (PTMs) of proteins are central in any kind of cellular signaling. Modern mass spectrometry technologies enable comprehensive identification and quantification of various PTMs. Given the increased number and types of mapped protein modifications, a database is necessary that simultaneouly integrates and compares site-specific information for different PTMs, especially in plants for which the available PTM data are poorly catalogued. Here, we present the Plant PTM Viewer (http://www.psb.ugent.be/PlantPTMViewer), an integrative PTM resource that comprises approximately 200,000 PTM sites for 17 types of protein modifications in plant proteins from five different species. The Plant PTM Viewer provides the user with a protein sequence overview in which the experimentally evidenced PTMs are highlighted together with functional protein domains or active site residues. The PTM sequence search tool can query PTM combinations in specific protein sequences, whereas the PTM BLAST tool searches for modified protein sequences to detect conserved PTMs in homologous sequences. Taken together, these tools facilitate to assume the role and potential interplay of PTMs in specific proteins or within a broader systems biology context. The Plant PTM Viewer is an open repository that allows submission of mass spectrometry-based PTM data to remain at pace with future PTM plant studies.

## INTRODUCTION

Posttranslational modifications (PTMs) lead to chemically different forms of proteins, so-called proteoforms (Smith *et al.,* 2013). Various proteoforms may be engaged in different protein complexes (Zhang *et al.,* 2016) or have altered stabilities (Nelson and Millar, 2015), structural conformations (Cho *et al.,* 2011) and activities (Huang *et al.,* 2018a). PTMs can also occur in a coordinated manner, resulting in a complex and interconnected landscape of protein regulation (Hunter, 2007, Minguez *et al.,* 2012). Although over 400 PTMs have been reported, only a subset of PTMs has been comprehensively characterized on a proteome-wide level by means of mass spectrometry (MS) due to the missing tools needed for their enrichment. On the other hand, due to technological advances in high-precision MS, up to thousands of PTM sites can now be identified in a single proteomics experiment (Choudhary and Mann 2010).

Our labs use proteomic tools to study PTMs occurring on plant proteins under various biotic and abiotic conditions (Jacques *et al.,* 2015, Walton *et al.,* 2016, Vu *et al.,* 2018). Thus far, protein phosphorylation is the best-characterized PTM in plants. For example, for *Arabidopsis thaliana* (Arabidopsis), approximately 57,000 phosphorylation sites are available in PhosPhAt 4.0 (Heazlewood *et al.,* 2008). Besides phosphorylation, recently reported PTMs include N-terminal myristoylation (Majeran *et al.,* 2018) and *O*-linked N-acetylglucosaminylation (*O*-GlcNAcylation) (Xu *et al.,* 2017) in Arabidopsis, and lysine 2-hydroxyisobuturylation (Meng *et al.,* 2017) and malonylation (Mujahid *et al.,* 2018) in rice (*Oryza sativa* spp. *japonica*). In addition to the identification of PTMs, several studies assessed differences in overall PTM levels, for instance under stress conditions (Jacques *et al.,* 2015, Vu *et al.,* 2018) or in developmental processes (Wang *et al.,* 2017). However, no integrative plant databases exist that capture such recently characterized modifications or their dynamics. Other plant protein modifications that have been identified by MS but rarely catalogued include cysteine oxidation and proteolytic processing. Oxidized cysteine sites have been detected in Arabidopsis and *Chlamydomonas reinhardtii*, following differential labeling of oxidized and reduced thiols (Liu *et al.,* 2014, Slade *et al.,* 2015). Proteolytically processed sites have been detected in Arabidopsis by enrichment of newly exposed protein N-termini via positional proteomics (Kleifeld *et al.,* 2010, Venne *et al.,* 2015), identifying substrates of the protease METACASPASE9 (Tsiatsiani *et al.,* 2013), processing associated with chloroplast protein import (Köhler *et al.,* 2015, Rowland *et al.,* 2015) and N-end rule substrates (Zhang *et al.,* 2015, 2018).

Interestingly, the increasing numbers of different PTMs further open up exciting possibilities to discover PTM interplays. Here, PTM interplay refers to an initial PTM that can act as a signal and influence the occurrence of another PTM (Hunter, 2007). Given the high quantity of identified PTMs, (species-specific) databases compiling PTM data for further exploration are needed. dbPTM (Lee *et al.,* 2006), PTMcode (Minguez *et al.,* 2015) and iPTMnet (Huang *et al.,* 2018b). As such databases often integrate various PTM types, they should facilitate the exploration of PTM interplays, leading to new hypotheses and experiments. For instance, incentivized by the overlap of protease cleavage sites and phosphorylation sites, human caspases were shown to cleave after phosphorylated serines (Seaman *et al.,* 2016). In plants, PTM interplay has been described as well. Based on observations in plant nitrite reductase and pyruvate dehydrogenase, nearby phosphorylation is inhibited by methionine oxidation that can act as a phosphorylation-regulating redox switch (Hardin *et al.,* 2009, Miernyk *et al.,* 2009). In addition, one or several phosphorylated residues have been shown to induce ubiquitination and subsequent protein degradation in a *cis*- regulatory manner (Hardtke *et al.,* 2000).

Given the vast amount of characterized PTMs on plant proteins, an enormous challenge now posed is on their functional characterization. Recently, a community effort has been suggested that would collect and assemble all PTM data for plant proteomes to systematically determine possible functional implications for plant proteins and the biological processes they are involved in (Friso and van Wijk, 2015). Although some PTM databases (Minguez *et al.,* 2015, Huang *et al.,* 2016, 2018b) provide plant protein modifications, PTM data are still largely uncatalogued and hardly explored in an integrative manner. To address this issue, we developed the Plant PTM Viewer (http://www.psb.ugent.be/PlantPTMViewer). In addition to ‘site-seeing’ of PTMs, we present several bioinformatics tools that help discover the hypothetical roles of plant PTMs in a protein-centric or systematic manner. Moreover, we attempted to visualize the quantitative differences in PTM levels between conditions. Hence, the Plant PTM Viewer offers the necessary framework to decode further plant PTM interplays to steer future plant protein studies.

## RESULTS

### A central resource for the exploration of PTMs of plant proteins

Plant PTM data were collected from 126 publications, often from supplementary tables. Approximately 200,000 experimentally evidenced PTM sites were mapped on plant proteins of five plant species, outcompeting the amount stored in other PTM databases (Figure 1; Table S1). Of the 17 PTM types reported in plants (Table S2), several were not catalogued for plants in any other PTM database. Additional information and statistics per PTM type are available via the ‘PTM Info’ tab. Besides the option to browse for PTMs per protein (‘Protein Search’ tab), the aim of the Plant PTM Viewer is to find the hypothetical role of a given protein. To this end, we introduced several tools that enable the functional dissemination of PTMs, including the possibility to search for user-defined PTM motifs (‘PTM Search’ tab) and to retrieve conserved PTMs in homologous proteins or protein regions (‘PTM BLAST’ tab). Please note that for each of the tools, a tutorial is available on the Plant PTM Viewer website (http://https://www.psb.ugent.be/PlantPTMViewer/).

**Figure 1.**
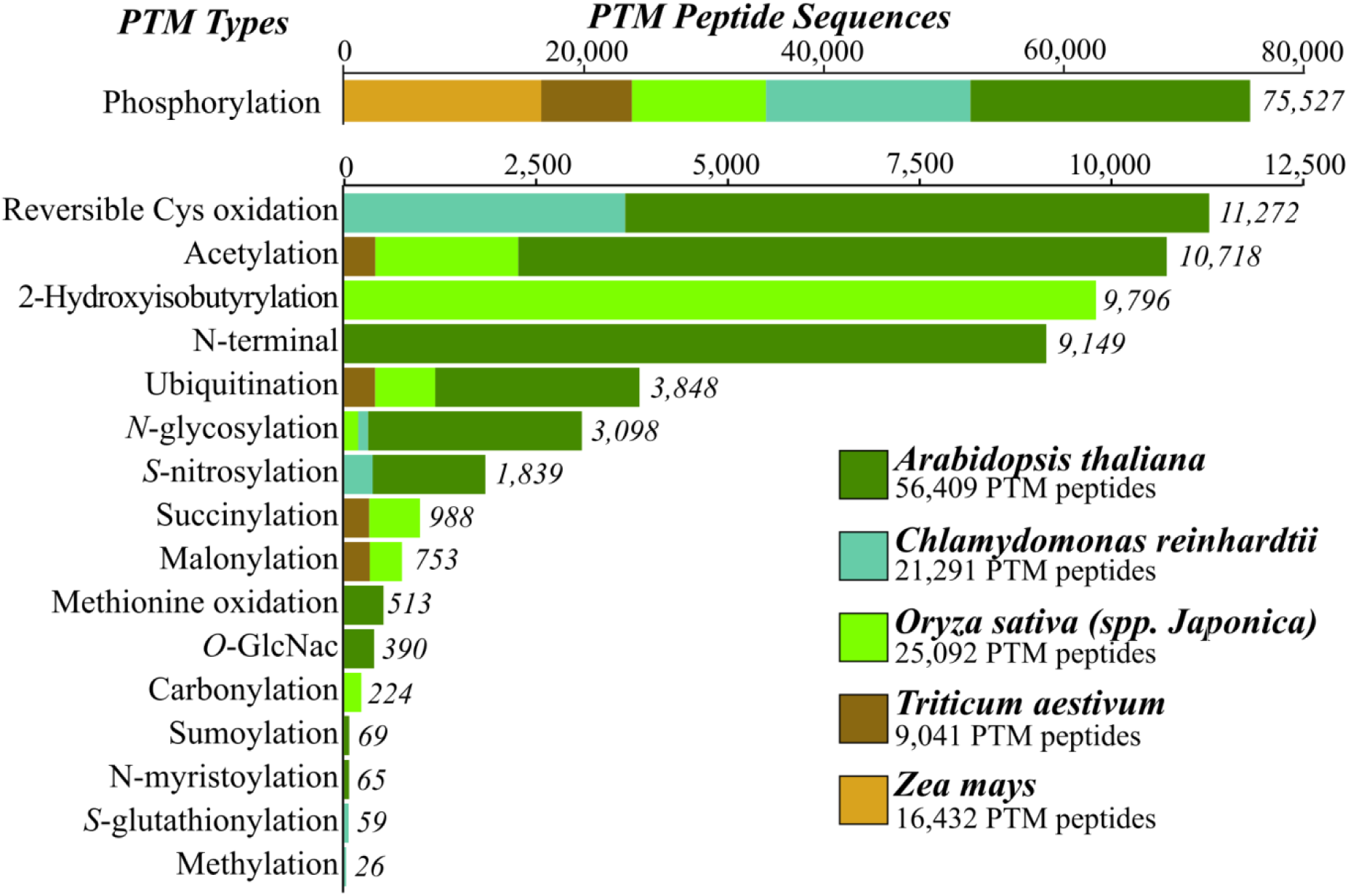
PTM Viewer datasets. Collected PTM peptide sequences identified by mass spectrometry for five plant species. All PTM peptides were displayed (x-axis) according to the PTM type (17 types, y-axis). Distinct peptide sequences per PTM type were counted. For phosphorylation, the scale was approximately 6.5-fold greater than for other PTM types. Bars were colored according to species. For more information regarding the PTM types, the reader is referred to the ‘PTM Info’ tab online.

### Examination of the complex PTM landscape for a protein of interest

With the ‘Protein Search’ tab a protein of interest can be searched, either via a protein identifier, a protein description, or a protein-matching amino acid sequence. The Plant PTM Viewer returns a protein-centric overview of all available PTMs in a tabular format, *i.e.* the ‘PTM table’, and a protein sequence overview in which all PTM sites are highlighted. As a test case, data are presented for ubiquitination, a highly conserved protein-based modification in which the small protein ubiquitin is covalently attached to substrate proteins often targeting them for degradation. Intriguingly, ubiquitin itself can be subjected to PTMs. For instance, in human ubiquitin, besides the polyubiquitin chain formation, substrate-bound ubiquitin can be acetylated, phosphorylated and modified by other ubiquitin-like molecules, referred to as the ubiquitin code (Swatek and Komander, 2016). To view the PTMs on plant ubiquitin and to explore possible parallels to the human ubiquitin code, we examined the highly conserved 76-amino-acid ubiquitin protein sequence in the ‘Protein Search’ tab. Several ubiquitin-encoding plant genes were found from which the protein sequence overview of the 76-amino-acid-long ubiquitin protein domain was extracted (Figure 2). In addition, the details for few modification sites described below are shown in the PTM table (Figure 2A, bottom). In the provided protein sequence overview the different experimentally evidenced ubiquitin modifications are shown, lysine ubiquitination being a prominent modification (Figure 2A). For the Arabidopsis and rice ubiquitin, all seven lysines were ubiquitinated, while only five lysines for wheat (*Triticum aestivum*). In the PTM table, every modified amino acid site and the respectively identified peptide sequence(s) can be consulted per individual study/experiment. For instance, in Arabidopsis, Lys48 ubiquitination was reported in three independent studies (Figure 2A, PTM table), giving some idea about the general prevalence of a modification. Besides polyubiquitination, various PTMs were identified on plant ubiquitins. Lys6 and Lys48 were found to be acetylated in Arabidopsis ubiquitin. These PTMs were partially conserved in wheat and rice, in which ubiquitin lysines were also often targeted by PTMs metabolically related to acetylation such as 2-hydroxyisobutyrylation (bu), succinylation (su) and malonylation (ma). So far, these latter modifications have not been characterized yet for Arabidopsis. For instance, in rice, all ubiquitin lysines, except Lys29, were targeted by either ma, bu and/or su (Figure 2A). Notably, the same six lysines were identified as being acetylated in human (Swatek and Komander, 2016). Interestingly, acetylation at Lys6 or Lys48 inhibited polyubiquitination in human, thereby shaping the ubiquitin branches (Ohtake *et al.,* 2015). Thus, we hypothesize that ubiquitin lysine acetylation, or metabolically related PTMs, might influence the type of ubiquitination also in plant proteins.

**Figure 2.**
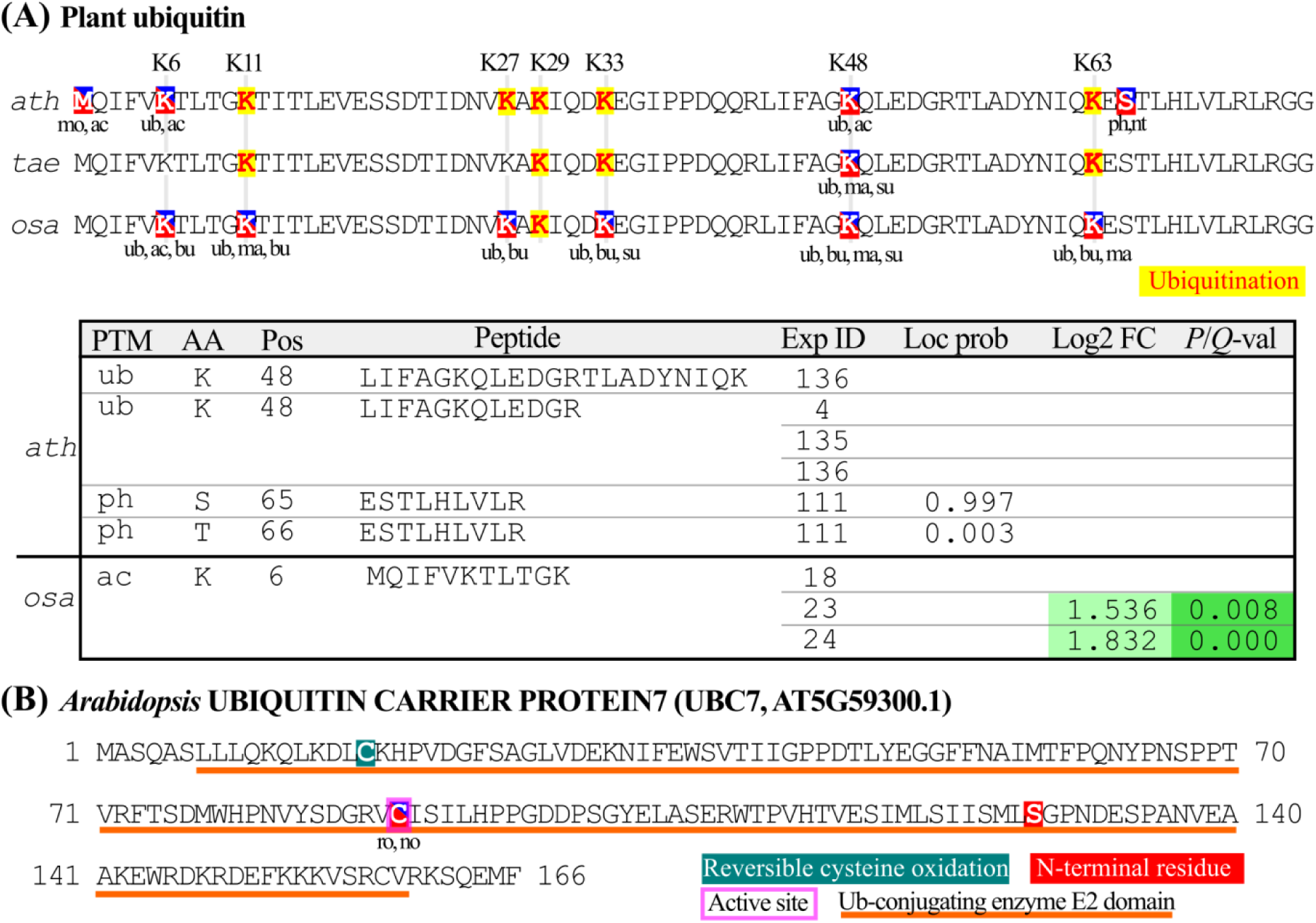
PTM protein overviews. (A) PTM protein sequence overview (top) and PTM table (bottom) of the ubiquitin protein domain encoded by Arabidopsis (*ath*) polyubiquitin 10 (AT4G05320), wheat (*tae*) TRIAE_CS42_6BS_TGACv1_515033_AA1666790, or rice (*osa*) LOC_Os06g46770. When the amino acid residues were subject to multiple modifications in the protein sequence overview, the two-letter code abbreviations of these modifications (Table S2) are indicated below the protein sequence. Ubiquitin lysines were specified with their respective position (K6, K11, K27, K29, K33, K48, and K63). Modification sites described in the text for Arabidopsis and rice ubiquitins are shown in the PTM table. For detailed information regarding the PTM table, the reader is referred to the tutorial and the respective protein overview pages online. (B) PTM protein sequence overview of Arabidopsis UBIQUITIN CARRIER PROTEIN 7 (UBC7, AT5G59300.1). The active site residue is indicated in a purple rectangle and the protein domain is underlined in orange. Abbreviations: Ac, acetylation; Bu, 2-hydroxyisobutyrylation; Ma, malonylation; Mo, methionine oxidation; No, S-nitrosylation; Nt, N-terminal; Ph, phosphorylation; Ro, reversible cysteine oxidation; Su, succinylation; Ub, ubiquitination.

Besides information on peptides and PTMs, the ‘PTM table’ provides additional peptide-level metadata that are sometimes reported in MS studies (Figure 2A, bottom). The Plant PTM viewer can store differential peptide abundancies between conditions, when the basic statistics are available (*e.g.*, log2-fold changes and *P*- or *Q*-values). In such cases, a comparison is considered as a different experiment, enabling visualization of PTM variabilities in a heat map-like display (Figure 2) and swift assessment of PTM intensities between conditions. For instance, differential lysine acetylation was assessed during early seed development in rice (Wang *et al.,* 2017). Here, Lys6 ubiquitin acetylation significantly increased after 3 and 7 days of seed development (Figure 2A, experiment ID 23 and 24, respectively). Interestingly, several ubiquitin-modifying enzymes have been implicated in seed size regulation of Arabidopsis, rice and wheat (Li and Li, 2014). Hence, differential ubiquitin acetylation might contribute to the seed development regulation. Besides differential abundancies, the PTMs identified by MS/MS spectra are often accompanied by localization probabilities within the identified peptide sequence. For instance, ubiquitin phosphorylation was assigned to Ser65 with a probability of 0.997 in Arabidopsis by means of the PhosphoRS algorithm (Taus *et al.,* 2011; Roitinger *et al.,* 2015). In human cells, this serine was shown to be phosphorylated by PINK1, leading to Parkin activation (Gladkova *et al.,* 2018). As Ser65 phosphorylation generates a structurally and functionally altered ubiquitin that can act as an independent signal (Wauer *et al.,* 2015), it is intriguing in the plant ubiquitin protein. In summary, by combining PTM data from diverse plant proteomics studies, a putative ubiquitin code in plants could easily be sketched, revealing analogy to the human ubiquitin code. However, the potential of the Plant PTM Viewer goes beyond ubiquitin, as the modification landscape of any plant protein of interest can be rapidly explored.

Another tool that facilitates hypothesis generation is the visualization of PTMs in a protein sequence view alongside their InterPro protein domains (Finn *et al.,* 2017) and sites that are annotated in UniProt Knowledgebase (UniProtKB) (UniProt Consortium, 2018) as binding or catalytic sites. For modifying enzymes controlling protein ubiquitination, such active domains and sites are well annotated. Ubiquitin is typically attached to substrate lysine residues through a three-step enzymatic process, involving E1 ubiquitin-activating, E2 ubiquitin-conjugating and E3 ubiquitin-ligating enzymes (Swatek and Komander, 2016). These enzymes have an active cysteine site that forms a thioester intermediate with ubiquitin. Here, the ubiquitin E2-conjugating enzyme UBIQUITIN CARRIER PROTEIN7 (UBC7, AT5G59300) (Figure 2B) was considered. Its E2 ubiquitin-conjugating protein domain (IPR000608) is posttranslationally modified by reversible cysteine oxidation (Cys17 and Cys89), S-nitrosylation (No, Cys88) and proteolytic processing (between Leu127 and Ser128). UniProtKB indicates that the redox-sensitive Cys89 is the active site involved in glycyl thioester intermediate formation. In Plant PTM Viewer, peptides ambiguously matching proteins encoded by different genes are flagged. The peptide ‘VCISILHPPGDDPSGYELASER’, identified by two independent experiments (Hu *et al.,* 2015; Liu *et al.,* 2015), is such an example, because it matches the active sites of both UBC7 and UBC13 (AT3G46460) that are closely related homologs. Thus, UBC7 (and/or UBC13) might be under redox control and be (partly) inactivated under oxidizing conditions, because the ubiquitin enzymes have been shown to be inactivated upon oxidation of the catalytic cysteine (Jahngen-Hodge *et al.,* 1997; Doris *et al.,* 2012). This example reveals that the Plant PTM Viewer steers the functional interpretation of PTMs in relation to annotated regulatory regions of a protein, thereby paving the road from PTM site-seeing to protein function.

### Assessment of the PTM interplay with the “PTM search tool”

The PTM search tool allows to search for ambiguous or specific amino acid motifs that are modified by one or more PTMs (for details, see help section or tutorial). This search ambiguity can be fine-tuned by users to address specific or more system-wide research questions. Searches can be restricted to retrieval of PTMs residing in known enzyme recognition sites (such as for kinases or proteases). For instance, substrate specificity was profiled for the Arabidopsis MITOGEN-ACTIVATED KINASE 3 and 6 (MPK3/MPK6) by means of a synthetic peptide library, delivering a Leu/Pro-X-pSer-Pro-Arg/Lys consensus motif for MPK6 (Sörensson *et al.,* 2012). Examination of this consensus motif in Arabidopsis proteins by ‘[LP]XS(ph)P[RK]’ with the PTM search tool returned 184 phosphosites uniquely matching 166 proteins (Figure 3, left panel), including the AT1G80180.1 protein that had been experimentally verified to be a MPK3 and MPK6 substrate (Sorensson *et al.,* 2012). In addition, combinations of PTMs can be searched to discover potential PTM interplays. Here, the possible interplay between phosphorylation and ubiquitination was considered, in which phosphorylated residues could inhibit or promote ubiquitination and act as so-called phosphodegrons (Filipčik *et al.,* 2017). To this end, we looked for ‘X(Ph)X{0,4}K(Ub)’, thus seeking phosphorylation of any residue (‘X(Ph)’), followed by lysine ubiquitination (‘K(Ub)’), spaced by none or maximal four amino acids (‘X{0,4}’). We also queried the reversed scenario, namely ‘K(Ub)X{0,4}X(Ph)’, in which a ubiquitinated residue is N-terminal to a phosphorylated one. Together these two queries returned 292 hits in 185 proteins (Figure 3, middle panel). Thus, such protein regions might hold putative phosphodegrons that promote or inhibit subsequent ubiquitination. As such, the PTM search tool can propose specific candidates and sites that might be of interest for future experimental validation. Moreover, the search results can optionally be restricted to match specified InterPro domains or UniProtKB-annotated protein sites.

**Figure 3.**
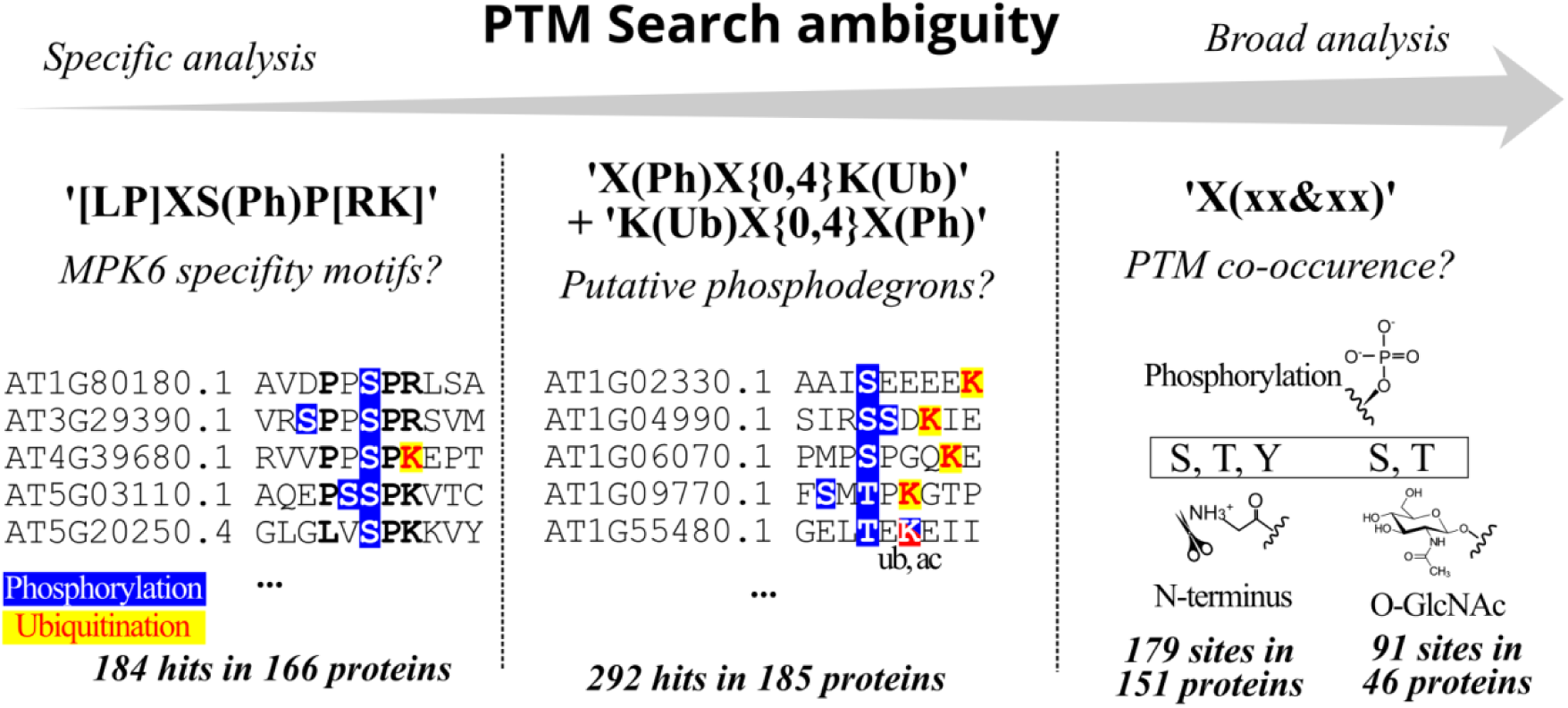
PTM Search with three queries and increased sequence and/or PTM ambiguity. For the MPK6 specificity and phosphodegron searches (left and middle panel), PTMs were highlighted in the returned hits. For ‘[LP]XS(Ph)P[RK]’, the consensus motif is in bold in the returned hits. For the search of the co-occurrence ‘X(xx&xx)’ (right panel), the co-occurrence of phosphorylation with N-terminal residues exposed after proteolytic processing (179 sites) and *O*-GlcNAc (91 sites) is shown. These PTM searches can be reproduced online by selecting *Arabidopsis thaliana* as species and unique peptides as advanced options. Additional information regarding the PTM search tool is provided in the PTM Search help section and in the tutorial.

The co-occurrence of different PTMs on the same amino acid is the most obvious form of PTM interplay, hinting at competition between modifying enzymes for a given amino acid. Such co-occurrence of PTMs can be explored with the term ‘X(xx&xx)’, i.e. looking for any amino acid (‘X’) targeted by at least two different modifications (‘xx&xx’). As an example, we focused again on phosphorylation. Phosphosites overlapped with 91 *O*-GlcNAc sites in 46 proteins and, surprisingly, also with 179 N-terminal residues exposed after proteolytic processing in 151 proteins (Figure 3, right panel). Further inspection of these phosphorylated sites revealed that 54 of these phosphosites resided at Thr2 or Ser2, thus at protein N-termini undergoing N-terminal methionine excision. Such phosphorylation at protein N-terminal residues had been reported to occur in photosynthetic membranes (Vener *et al.,* 2001). In some cases, the N-terminal modification status was even more complex. Ser2 of the NUCLEOSOME ASSEMBLY PROTEIN1 (NAP1, AT2G19480.1) could be exposed by N-terminal methionine excision and be N-terminally acetylated (Zhang *et al.,* 2015, 2018), also be phosphorylated (Roitinger *et al.,* 2015) and even N-terminally ubiquitinated (Walton *et al.,* 2016). Hence, the N-terminal modification landscape of some proteins might be more complex and dynamic than anticipated.

### Discovery of conserved PTMs in proteins within or across species with ‘PTM BLAST’

Whereas the PTM sequence search tool can return specific PTM patterns in proteins, the ‘PTM BLAST’ tool can identify similar protein regions with conserved PTMs within or across species. ‘PTM BLAST’ can use a (modified) protein sequence as query for a default protein BLAST to all the proteins in the Plant PTM Viewer (Figure 4A). In a next step, the obtained aligned protein sequences are cross-checked for alignment of PTMs of differing (‘align’) or identical (‘match’) types. The top results are sorted by the numbers of PTM alignments (Figure 4A), allowing the swift identification of conserved PTMs within or across plant species. BLAST alignments can be visualized and their respective PTMs are indicated. Ambiguous peptide sequences matching both aligned proteins can be omitted. Such peptides can appear in protein alignments within species due to ambiguous peptide-to-protein assignment. ‘PTM BLAST’ can be launched from each PTM Viewer protein overview, querying the entire protein sequence or a region thereof, thus restricting the BLAST search to a region of interest, for instance a protein domain.

**Figure 4.**
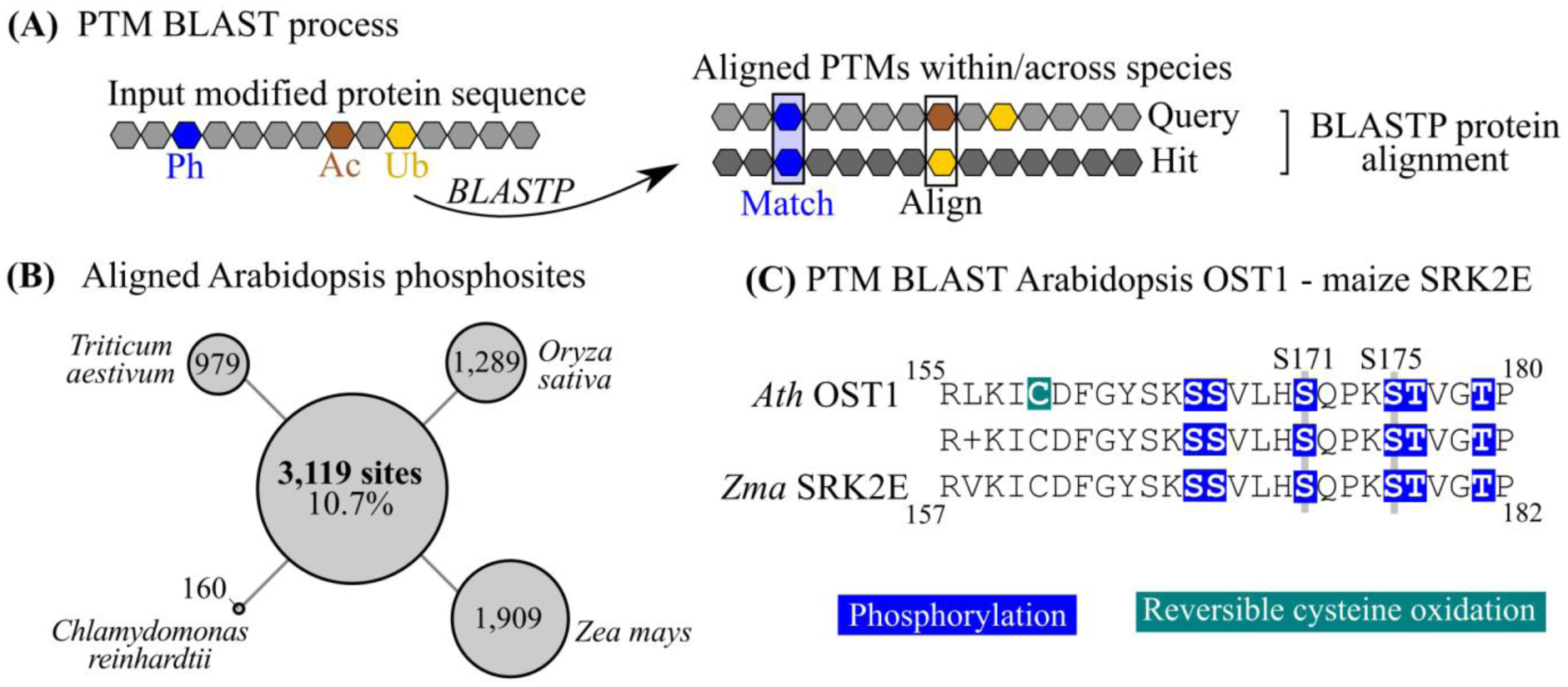
‘PTM BLAST’ and sequence alignment. (A) A default protein BLAST (BLASTP) carried out by ‘PTM BLAST’ for a given modified protein sequence, *e.g*., phosphorylation (Ph, blue), acetylation (Ac, brown) and ubiquitination (Ub, yellow). Protein alignments of significant BLASTP hits can be visualized with PTMs highlighted. Two PTM alignment cases are counted: the alignment of modified residues regardless of the PTM type (‘align’ type; *e.g.* Ac and Ub) and of the same PTM type (‘match’ type). (B) Alignment of Arabidopsis phosphorylation sites in orthologous proteins. The number of phosphorylation sites that aligned exactly to a phosphosite in an orthologous protein (BLASTP E-value < 0.5, alignment length > 80% longest sequence and amino acid identity > 60%) was counted. (C) ‘PTM BLAST’ alignment of conserved phosphosites, including Ser171 (S171) and Ser175 (S175), between Arabidopsis OPEN STOMATA1 (OST1, AT4G33950.1) and its maize ortholog SRK2E (Zm00001d033339_P002).

To evaluate the evolutionary conservation of Arabidopsis PTMs, we ran the ‘PTM BLAST’ for all canonical Arabidopsis proteins, keeping track of the number of Arabidopsis PTMs that aligned to other types of PTMs per species. For instance, in case of phosphorylation that is the most abundant PTM type represented in all species (Figure 1), 3,119 phosphosites in Arabidopsis (10.7% of the searched sites) aligned exactly to phosphosites of any other plant species (Figure 4B). Moreover, 69 sites aligned exactly in all four species, corresponding to 43% of the available *Chlamydomonas* phosphosites. Hence, a subset of phosphosites in cyanobacteria seems relatively well conserved throughout the plant clades. Many of these 69 plant ultra-conserved phosphosites map to kinases, such as MPK1, MPK2, MPK3, MPK4, MPK6, MPK7 and MPK11. Another example is OPEN STOMATA 1 (OST1, AT4G33950), in which both Ser171 and Ser175 are conserved phosphosites that reside in the OST1 activation loop of all species. Noteworthy, in Arabidopsis, both serines were found to be phosphorylated after hyperosmotic or abscisic acid treatment and Ser175 to be crucial for the OST1 phosphorylation activity (Belin *et al.,* 2006). As an example of a ‘PTM BLAST’ alignment, we present a protein region alignment between the Arabidopsis OST1 and its maize (*Zea mays*) ortholog SRK2E (UniProt accession B4FQ40) that, besides Ser171 and Ser175, has four supplementary conserved phosphosites (Figure 4C).

Besides querying a PTM Viewer protein, any custom protein sequence, with or without specified PTMs included in the Plant PTM Viewer (Table S2), can be searched via ‘PTM BLAST’. As a case example, we selected the human glyceraldehyde-3-phosphate dehydrogenase (GAPDH), because it is an essential housekeeping gene for which a rich PTM diversity is evident, as substantiated by the 25 sites of five PTM types reported in UniProtKB that are included in the PTM Viewer (UniProtKB P04406) (Figure S1A). Therefore, the complex PTM landscape of the human GAPDH was used as input for the ‘PTM BLAST’, resulting in exact PTM alignments for the GAPDH orthologs of all the five plant species. For GAPDH C2 (GAPC2) of Arabidopsis, 10 conserved PTMs were retrieved, including seven phosphorylation sites, two acetylation sites and a S-nitrosylation site (Figure S1B). In addition, also malonylated sites of the human GAPDH, Lys194 and Lys216, were conserved in the wheat and rice GAPDH orthologs. Thus, ‘PTM BLAST’ can return aligned PTMs for a given, non-plant input sequence as illustrated here for the human GAPDH.

## DISCUSSION AND PERSPECTIVES

The Plant PTM Viewer enables researchers to consult the PTM landscape for plant proteins in a user-friendly manner. The current collection of plant PTM data entails approximately 200,000 sites of 17 PTM types of five plant species. The Plant PTM Viewer offers various tools that facilitate the functional interpretation and hypothesis formulation of PTMs. In contrast to other PTM databases, the ‘PTM search’ tool allows the search for combinations of PTMs or of single PTMs in user-defined amino acid motifs. Thanks to an easy fine-tuning, the biological research question at hand can be addressed and putative protein PTM candidates can be hypothesized. In addition, the ‘PTM BLAST’ tool can reveal aligned PTMs across or within species, thereby enhancing the confidence and relevance of these PTMs. In the future, these tools could be extended with, for instance, assessment and visualization of overrepresented gene sets and sequence motifs in the case of a PTM search, or multiple sequence alignments of homologous protein regions delivered by ‘PTM BLAST’. Like the database information, all the results of the Plant PTM Viewer can easily be exported, facilitating the plant PTM research by directing both future *in silico* systematic analyses and wet-lab experiments.

The Plant PTM Viewer strives to be and remain a central resource for plant researchers. To this end, we strongly encourage the plant science community to actively contribute to the Plant PTM Viewer by submitting peer-reviewed PTM data. The solid Plant PTM Viewer framework for protein visualization and tools can be easily expanded to deal with additional PTM sites, including those from newly characterized PTM types and additional plant species. Despite the phosphorylation dominance in the PTM research, we expect extra PTM types to be better characterized in the future. However, due to the increasing wealth of reported PTMs in proteins, an important future challenge will be to determine whether PTMs are functionally important for protein functions or not. In this respect, acquisition of quantitative information will become increasingly important to determine abundant PTMs or those that differ between conditions. Although differential PTM data are still in their infancy, we feel that we have taken an initial step toward a differential PTM browser, similar to GENEVESTIGATOR for differential gene expression (Hruz *et al.,* 2008).

## EXPERIMENTAL PROCEDURES

### PTM data and metadata collection and processing

From 126 peer-reviewed MS studies on plant proteomes, PTMs with their respective peptide sequences were collected, including the phosphopeptide data extracted from the PhosPhAt 4.0 database (Heazlewood *et al.,* 2008). In total, 128,265 peptide sequences were assembled that reported PTMs from five species: Arabidopsis, maize, common wheat, rice and *Chlamydomonas reinhardtii* (Figure 1). Peptide sequences carrying PTMs were reallocated to all proteins (i.e., including splice forms) from the latest reference proteomes of these five species (Table S3), resulting in approximately 200,000 PTM sites in representative proteins. Ambiguous protein interference frequently arises in plants due to extensive duplication events (Vanneste *et al.*, 2014). Hence, in the Plant PTM Viewer, approximately 20% of the PTM sites matched to proteins encoded from different genes. To address this issue, ambiguous peptides that matched multiple proteins were flagged in the PTM Viewer. When differential abundance statistics between conditions were reported, the log2-fold change and significance values (*e.g., P*-value and false discovery rate) were stored as well, occurring currently for approximately 4% of the nonredundant list of PTM peptides. In addition, the PTM localization probabilities were saved when available, mostly provided by the MaxQuant PTM Score (Olsen *et al.,* 2006) or PhosphoRS (Taus *et al.,* 2011) for the majority (72%) of the PTM sites, due to the high number of phosphorylation studies that use localization probability algorithms.

The PTM data were accompanied with various protein and experimental metadata. Protein identifiers were cross-referenced with UniProtKB (UniProt Consortium, 2018) and the plant comparative genomics platform PLAZA (Van Bel *et al.,* 2018) for identical protein sequences. For the original MS studies, the respective PubMed identifier and the PRIDE (Vizcaíno *et al.,* 2016) accession data of the raw MS data, if deposited, were stored. In addition, relevant experimental information, such as plant tissue and genotype, stress condition, PTM enrichment methodology and tandem MS search parameters were retained as well.

### System configuration

The Plant PTM Viewer is accessible via http://www.psb.ugent.be/PlantPTMViewer. A web interface for data browsing, searching and displaying was implemented in PHP. To enable searching for PTMs, sequence-based queries are translated to specific regular expressions by the use of various wildcards and operators for PTMs. For instance, ‘X’ can be used to denote any amino acid and ‘[ST]’ can specify either serine or threonine. Hence, the retrieval of complex patterns is allowed, whereby amino acids and PTM types can be specified in a flexible manner. Screen resolution is auto-adaptable for user-friendly browsing from wide screens to smart phones. The Plant PTM Viewer can be accessed programmatically. The Plant PTM viewer data are maintained in a MySQL relational database, of which an overview is given in Figure S2. For enhancement of the functional interpretation, the Plant PTM Viewer is connected to the InterPro domain database (Finn *et al.,* 2017), as well as to UniProtKB (UniProt Consortium, 2018).

## ACKNOWLEDGMENTS

The authors thank Lam Dai Vu, Tingting Zhu, Natalia Nikonorova, Silke Jacques, Alan Walton, Lukas Braem, and the plant research community characterizing PTMs in plants for providing the necessary input for the Plant PTM Viewer. In addition, they acknowledge Michiel Van Bel and Frederik De Laere for providing IT support, Simon Stael and Kai Xun Chan for critical reading and comments, and Martine De Cock for help with the manuscript preparation. This work was supported by grants from the Research Foundation-Flanders (grant number G.0038.09N [to K.G. and F.V.B.] and Excellence of Science project no. 30829584 [to F.V.B.]), the Ghent University Multidisciplinary Research Partnership [“Ghent BioEconomy” (Project 01MRB 510W) to F.V.B.] and the Ghent University Special Research Fund (BOF 01J11311 [to F.V.B.]).

## SUPPORTING INFORMATION

Additional Supporting Information may be found in the online version of this article.

**Figure S1.** ‘PTM BLAST’ of the human glyceraldehyde-3-phosphate dehydrogenase (GAPDH).

**Figure S2.** The PTM Viewer MySQL database scheme.

**Table S1.** Comparison of the PTM Viewer unique PTM peptides per species versus PTM sites stored in other PTM databases.

**Table S2.** Seventeen plant PTMs included in PTM Viewer

**Table S3.** Species reference annotations.

